# Early warning indicators for heat-induced mortality in temperate tree saplings

**DOI:** 10.64898/2026.08.18.745401

**Authors:** Clara Stock, Stefanie Dumberger, Mirjam Meischner, Melissa Wannenmacher, Kathrin Kühnhammer, Jürgen Kreuzwieser, Simon Haberstroh, Christiane Werner

**Affiliations:** Ecosystem Physiology, Faculty of Environment and Natural Resources, University of Freiburg, Freiburg, Germany; Center for Volatile Interactions (VOLT), Department of Biology, University of Copenhagen, 2100 Copenhagen, Denmark

**Keywords:** tree mortality, heat stress, early warning indicators, legacy effects, *Fagus sylvatica*, *Pseudotsuga menziesii*, *Picea abies*

## Abstract

- Globally, forest ecosystems face widespread mortality events. However, the independent impacts of distinct stressors, such as heat stress vs edaphic drought, remain poorly understood and physiological early warning indicators for tree mortality are urgently required.
- We exposed well-watered saplings of *Fagus sylvatica*, *Pseudotsuga menziesii* and *Picea abies* to summer heat waves and subsequent natural winter-desiccation. Physiological parameters (e.g. gas exchange, water uptake velocity via ^2^H labelling, and volatile organic compound emissions) were monitored throughout the growing season and survival was assessed regularly until subsequent spring to capture immediate and delayed mortality as a consequence of legacy effects.
- Heat exposure without soil water deficit, followed by winter desiccation, triggered species-specific mortality rates (51.8% *F. sylvatica*, 48.2% *P. abies*, 16.9% *P. menziesii*), with *P. abies* exhibiting significantly faster mortality response than the other species. Reduced water uptake, lower stomatal conductance, impaired photosynthetic efficiency, and altered VOC emissions distinguished non-surviving from surviving saplings months before visible damage in all three species.
- Heat stress drives mortality independent of edaphic drought, with sub-lethal physiological indicators detectable up to 10 months before visual signs. These early warning indicators could enable damage detection before lethal thresholds are crossed, offering new strategies for mitigating climate change-driven forest decline.

## Introduction

The global trend of increasing air temperature (IPCC 2023) is particularly pronounced on the European continent, where warming happens twice as fast as the global average (Chen 2026). During the last decades, this has manifested in record-breaking heatwaves across Europe, including those in 2003, 2018, 2022 and 2026 (Schuldt et al. 2020; Spiecker and Kahle 2023; Vautard et al. 2023; Chen 2026). These extreme events, often compounded by additional stressors such as edaphic drought, are increasingly driving widespread tree mortality and threatening the stability of European forest ecosystems (Allen et al. 2010; Schuldt et al. 2020; Haberstroh et al. 2022). Such impacts manifest both as immediate mortality and as legacy effects during the subsequent years (Yu et al. 2025; Pohl et al. 2023). Yet, understanding the mechanistic drivers of tree mortality and predicting these die-offs remain challenging (McDowell et al. 2008; Allen et al. 2010; Hammond et al. 2019). Key uncertainties persist regarding the physiological causes and timing of death, specific lethal thresholds, and intra-and interspecific differences in stress resilience (McDowell et al. 2008; Hammond et al. 2019). Resolving these questions is essential to improve mortality forecasts under a changing climate and will facilitate the identification of early warning indicators prior to crossing lethal thresholds (Trugman et al. 2021).

The primary mechanism driving tree mortality under drought is hydraulic failure, resulting from extensive xylem embolism during severe stress (McDowell et al. 2008; Anderegg et al. 2012; Adams et al. 2017). Additionally, prolonged stomatal closure can deplete non-structural carbohydrate reserves, leading to carbon starvation that may further increase mortality risk (Anderegg et al. 2012; Sevanto et al. 2014; Adams et al. 2017). Critical physiological thresholds are often exceeded long before catastrophic hydraulic failure or visible foliar damage occurs, complicating the early detection of potentially lethal damages (Adams et al. 2017; Blackman et al. 2019; Hammond et al. 2019). So far, most research has focused on drought-induced mortality (Choat et al. 2018; Arend et al. 2021; McDowell et al. 2022), whereas emerging evidence suggests that extreme heat and high vapor pressure deficit (VPD) can act as independent drivers of mortality without the co-occurrence of edaphic drought (Grossiord et al. 2020; Breshears et al. 2021; Marchin et al. 2022; Still et al. 2023; Werner et al. 2026). Disentangling the effects of these stressors, e.g. by analysing the independent impact of heat, is crucial for understanding the underlying mechanisms of mortality. For this, it is essential to assess both immediate and delayed mortality. Legacy effects leading to delayed mortality in previously stressed trees during subsequent years, potentially triggered by secondary stress events, have been reported in response to various stressors, such as drought, heat and post-stress insect or pathogen infestations (Trugman et al. 2018; Buras et al. 2020; Schuldt et al. 2020; Pohl et al. 2023). Critically, short term triggers, such as winter desiccation, limiting root water uptake, after the exposure to a long term stressor can exacerbate mortality effects (Peguero-Pina et al. 2011; Voltas et al. 2013). Such effects may be particularly pronounced following heat stress, as heat-induced damage often progresses gradually (Hüve et al. 2011; Birami et al. 2018). This underscores why long-term monitoring is essential to fully capture tree mortality dynamics and avoid underestimating long-term impacts.

Heat exposure affects plant physiology in multiple ways, including reduced net photosynthesis due to decreased Rubisco activase activity, inhibited electron transport capacity via decreased photosystem II (PSII) activity, increased mitochondrial and photorespiration, and compromised thylakoid membrane stability (Hüve et al. 2011; Teskey et al. 2015; Guha et al. 2018), amongst others. At the whole-tree level, heat stress reduces growth and leaf area, and, particularly when combined with edaphic drought, can lead to tree mortality (Teskey et al. 2015). Key protective mechanisms during heat exposure include transpirational cooling and the emission of volatile organic compounds (VOCs) (Peñuelas and Llusià 2003; Loreto and Schnitzler 2010; Hüve et al. 2011; Werner et al. 2020). In the present study, we analysed the angiosperm *Fagus sylvatica* L. and the two conifers *Pseudotsuga menziesii* (Mirb.) Franco and *Picea abies* (L.) H. Karst, representing important European tree species. These species vary significantly in their capacity for both protective mechanisms (Kleist et al. 2012; Teskey et al. 2015). Conifers typically exhibit more isohydric water-use strategies and greater hydraulic safety margins than angiosperms (Choat et al. 2012; Schumann et al. 2024; Paligi et al. 2025). Furthermore, conifers possess specialised VOC storage structures, whereas many angiosperms, including *F. sylvatica,* lack such storage capacity (Dindorf et al. 2006; Holzke et al. 2006; Ghirardo et al. 2010; Niinemets et al. 2011). The co-occurrence of edaphic drought intensifies heat stress effects (Birami et al. 2018; Haberstroh et al. 2026; Werner et al. 2026). Consequently, water availability during heat exposure is critical, as it facilitates survival through transpirational cooling preventing lethal overheating of the leaves (Bauweraerts et al. 2013; Ruehr et al. 2015; Teskey et al. 2015). However, increasing evidence suggests that extreme heat can cause whole-tree mortality even under well-watered conditions (Birami et al. 2018; Still et al. 2023). For instance, Birami et al. (2018) observed sapling mortality of *Pinus halepensis* under heat treatments, proposing heat-induced membrane damage and protein denaturation as the primary drivers of mortality rather than hydraulic failure. Furthermore, within-species variability in survival was explained by the individual capacity for transpirational cooling (Birami et al. 2018), suggesting that intraspecific differences in thermoregulation determine individual stress resilience.

Predicting tree mortality remains a key challenge across both inter- and intraspecific comparisons (McDowell et al. 2008; Hartmann et al. 2022; Mosig et al. 2026). Deciduous European beech (*Fagus sylvatica* L.) and coniferous Douglas fir (*Pseudotsuga menziesii* (Mirb.) Franco) and Norway spruce (*Picea abies* (L.) H. Karst) are among the most economically and ecologically important tree species in European forests (Spiecker et al. 2019; Obladen et al. 2021). Over the last decades, drastic die-backs have been reported particularly for *P. abies*, which is currently considered the most vulnerable conifer in European forests (Spiecker et al. 2019; Arend et al. 2021; Anders et al. 2025). Similarly, mortality rates in *F. sylvatica* and *P. menziesii* have increased in response to environmental stress, albeit to a lesser extent (Leuschner 2020; Schuldt et al. 2020; Obladen et al. 2021; Schumann et al. 2024; Leuschner and Meinzer 2024; Cavelier et al. 2025). Vulnerability to environmental stress is often attributed to divergent hydraulic strategies, such as differences in isohydricity and hydraulic safety margins (McDowell et al. 2008; Meinzer et al. 2009; Choat et al. 2012, 2018; Schumann et al. 2024; Paligi et al. 2025). However, these traits are not universal predictors of mortality, as additional factors such as site conditions, rooting depth or genetics also play critical roles (Schumann et al. 2024). At the intraspecific level, genetic variation affects traits such as vulnerability to xylem cavitation, water use efficiency and stomatal density and size and can significantly influence survival (McDowell et al. 2008). Accounting for this variation could therefore improve predictions of individual resilience (Teskey et al. 2015).

In this study, we exposed potted saplings of *F. sylvatica*, *P. menziesii* and *P. abies* to naturally occurring heat waves during the 2023 growing season. Despite regular watering and the absence of edaphic drought, we observed significant mortality across all species during the growing season. Subsequently, survivors were exposed to naturally occurring winter desiccation in early 2024 resulting in a legacy mortality event across all species. By assessing mortality rates and key physiological parameters, including gas exchange, leaf water potential, chlorophyll fluorescence and water uptake velocity assessed via ^2^H pulse labelling throughout the experiment we aimed to 1) assess species-specific mortality and legacy effects in response to heat exposure without concurrent drought, 2) analyse the underlying, physiological mechanisms driving heat-induced mortality and 3) identify early indicators that distinguish survivors from dying plants before lethal thresholds are crossed. Addressing these objectives, we hypothesise that 1) *P. menziesii* is more stress-resilient than *F. sylvatica* and *P. abies*, with *P. abies* exhibiting the highest susceptibility to heat-induced mortality, 2) heat exposure causes substantial damage to both the hydraulic and the photosynthetic system in vulnerable plants and 3) non-destructive physiological measurements can indicate potentially lethal physiological impairments before visual damage is detectable.

## Material and Methods

### Plant Material and Study Site

The study was performed on 2-4 year old saplings of *Fagus sylvatica* L. (European Beech), *Picea abies* (L.) H. Karst (Norway Spruce) and *Pseudotsuga menziesii* (Mirb.) Franco (Douglas fir) from tree nurseries located in southern Germany (Baumschule Haage GmbH & Co. KG, Leipheim, Germany and Gustav Burger Forstbaumschulen, Zell am Harmersbach, Germany). The saplings were potted between April and June 2023 in 38L Euroboxes (AUER GmbH, Amerang, Germany). Each box was equipped with two saplings in either heterospecific (*F. sylvatica* with *P. menziesii* or *F. sylvatica* with *P. abies*) or monospecific combinations (two individuals of each species). Whole-tree harvesting at the end of the experiment confirmed that root systems did not overlap between co-planted saplings. Given the absence of root competition, no interspecific interactions were expected. This was supported by the lack of significant differences among species combinations. Therefore, data were pooled across all combinations for subsequent analyses. Soil contained 20% peat and was mixed with sand (v:v, 5:1). The boxes were fertilized with 2g NPK slow-release fertilizer (Osmocote, 15 + 9 + 12 + 2MgO). From April 2023 until April 2024, the boxes were placed on a scaffold with transparent rain-out shelter on the campus of the University of Freiburg. Thus, plants were exposed to outside conditions with controlled irrigation. In total there were 278 plant individuals (110 *F. sylvatica*, 83 *P. menziesii* and 85 *P. abies*). Mortality screenings were performed regularly on all individuals. Regular physiological and VOC measurements were performed on randomly selected individuals (*F. sylvatica:* n=36; *P. menziesii:* n=24; *P. abies:* n=24).

### Meteorological and soil moisture measurements

The study site was equipped with a HOBO station (Onset Computer Corporation, Bourne, USA) measuring air temperature (T_air_), relative air humidity (S-THB-M002) and photosynthetic photon flux density (PPFD) (S-LIA-M003) every 5 minutes from July 2023 until April 2024. Out of the 60 boxes, which were used for the physiological measurements, 32 were equipped with a 5TM, EC-5 or GS1 soil moisture sensor (METER group, Munich, Germany), evenly distributed across the different species combinations. Soil moisture sensors were connected to EM50 loggers (METER group, Munich, Germany) and recorded values every 5 minutes from June 2023 until March 2024. Soil moisture measurements ended earlier, because they were needed in another experiment. Irrigation was adjusted to prevent water stress and maintain a target range of 20–30 % volumetric water content (VWC). The rain-out shelter altered T_air_ relative to ambient conditions, with daily mean differences averaging 0.87 ± 0.71 °C (range: -0.82 to 4.31 °C). This warming effect was most pronounced during sunny days, leading to higher maximum air temperatures, hence intensifying the heat waves (Fig. S1). Hot days, tropical nights and frost days under the rain-out shelter were recorded throughout the experiment, defined as days with maximum T_air_ > 30 °C, minimum T_air_ > 20 °C and minimum T_air_ < 0 °C, respectively (Crespi et al. 2020).

### Experimental design

From July 2023 until Mai 2024, plants were exposed to natural meteorological conditions beneath the rain-out shelter with irrigation being the only controlled environmental variable. Throughout the experiment, distinct physiological measurements and screenings were performed regularly: Mortality screenings and predawn measurements of leaf water potential (Ψ_PD_) and maximum quantum use efficiency (Fv/Fm) were assessed under natural conditions. Additionally, a climate chamber experiment with a ^2^H pulse labelling was conducted to investigate water uptake and transport velocities (Fig. 1). For this purpose, plants were temporarily transferred to walk-in climate chambers (ThermoTEC, Weilburg, Germany) during three periods throughout the experiment (22^th^ of July – 07^th^ of August; 16^th^ of August – 15^th^ of September; 4^th^ – 20^th^ of October). Inside the climate chambers, one branch per plant was enclosed in leaf cuvettes that were connected to an automated measurement system (Werner et al. 2020) enabling the on-line measurement of leaf gas exchange (i.e. net carbon assimilation and transpiration) and the sampling of VOCs (measurement system described in detail below). Plants were measured in seven consecutive charges, since the capacity of the automated system was limited. After minimum two days of acclimatization, each plant was analysed continuously for at least 24 hours. During the second climate chamber measurements, each box was irrigated with 500mL of ^2^H enriched water (δ^2^H_2_O: 5295‰ ± 64‰) and on-line measurements were performed up to six days during this period to ensure capturing the ^2^H water pulse in the transpiration. For each climate chamber period and tree individual, one VOC sampling for gas chromatography-mass spectrometry (GC-MS) analysis was performed and one branchlet was sampled for analysis of water-soluble organic matter (WSOM). The destructive sampling of branchlets was performed after the VOC measurements to avoid a burst of VOCs due to mechanical damage. Environmental parameters in the climate chambers were set to elevated air temperatures of 28 °C/18 °C T_air_ (day/night), 60% relative air humidity (day and night), 15h light, 7h dark, 60min dawn and dusk, each, and ∼600 µmol m^-2^ s^-1^ of PPFD for the first and second climate chamber measurement period. During the third measurement period in the climate chambers, parameters were adapted to later season, with 23 °C/13 °C T_air_ (day/night), 60% relative air humidity (day and night), 12h light, 10h dark, 60min dawn and dusk each and ∼500 µmol m^-2^ s^-1^ PPFD.

**Fig. 1.**
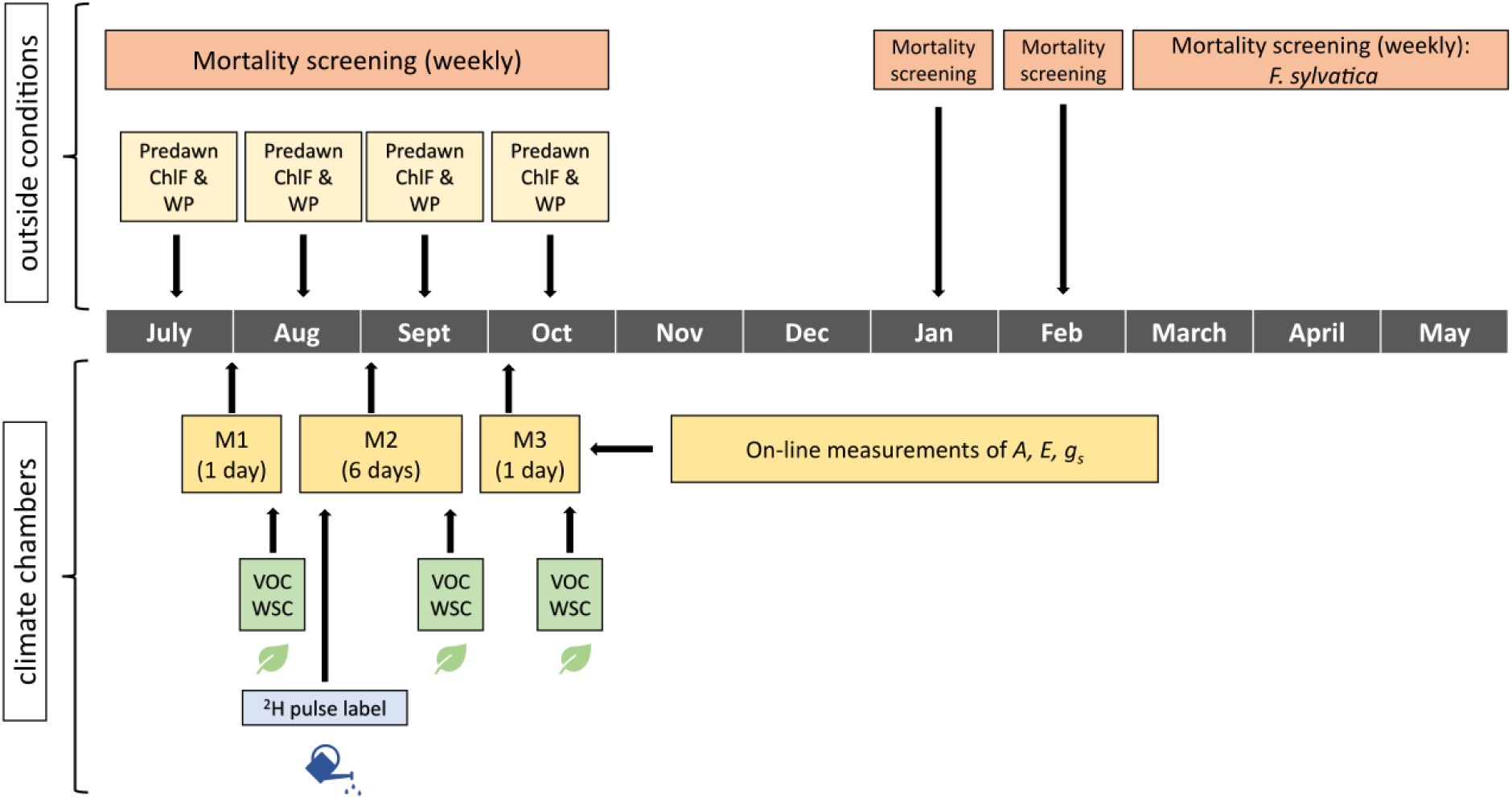
Schematic overview of the physiological measurements and screenings from July 2023 until May 2024. Mortality screenings, chlorophyll fluorescence (ChlF) and water potential (WP) measurements were performed under outside conditions. On-line measurements of gas exchange parameters (assimilation (*A*), transpiration (*E*) and stomatal conductance (*g_s_*)) and sampling of volatile organic compounds (VOCs) and water-soluble carbohydrates (WSOMs) were performed inside the climate chambers during three measurement periods (M1, M2, M3). 2H pulse labelling was applied at the beginning of the second climate chamber measurement period (M2).

### Mortality screening

From July until October 2023, mortality screenings were conducted every 1-2 weeks on all 278 saplings. Foliar damage was visually estimated for each individual. Scores ranged from 0 % (entirely green) to 100 % (complete foliar discoloration or defoliation). Some of the *F. sylvatica* exhibited reflushing, producing up to three successive leaf cohorts during the growing season, which resulted in fluctuations in damage scores. For all species, individuals reaching 100 % foliar damage without subsequent recovery were classified as dead. Time of death was defined as the moment when 100 % foliar damage was first reached. We note that the mortality timings in our study were determined based on visual observation, which has some inherent uncertainties on the precise dating of mortality. Determining the exact moment with non-destructive measurements remains a key challenge in tree mortality research (Hammond et al. 2019). In January and February 2024, additional screenings were carried out on all plants to evaluate legacy effects of mortality. Since leaf mortality could not be assessed on dormant leafless *F. sylvatica* saplings, weekly screenings were performed on *F. sylvatica* from 28^th^ of February to 21^st^ of May 2024, documenting bud-break. Individuals of *F. sylvatica* that showed bud activity during spring, but died before reaching full leaf-out were classified as legacy mortality plants. Based on the mortality screenings, plants were divided into three groups: plants that died during the growing season 2023 (mortality group), plants that died in 2024 (legacy group) and plants that survived (survivor group). Investigated tree individuals were not equally distributed across the three groups with the mortality group reaching especially low numbers of replicas due to ongoing mortality.

### Physiological measurements

Physiological measurements were conducted throughout the 2023 growing season on a subset of saplings. Individuals that died during 2023 were monitored until death, resulting in a progressive decline in replicate numbers for this category over time. Crucially, measurements were not performed during peak heat events but under moderate conditions (either in the climate chambers at T_air_ = 28/23 °C or at predawn outdoors). Consequently, the data reflect physiological performance following heat exposure rather than during acute stress. The final measurement campaign during October 2023 was conducted on all plants exhibiting no visible mortality symptoms, establishing a baseline comparison between potential survivors and the legacy mortality group prior to winter dormancy. Notably, the mortality group could not be included in this campaign, as individuals in this group had already died before the October measurement campaign. In October 2023, *F. sylvatica* saplings from legacy and survivor plants already showed signs of senescence.

#### Discrete leaf water potential and chlorophyll fluorescence measurements

From July until October 2023, predawn measurements of leaf water potential (Ψ_PD_) and chlorophyll fluorescence were performed once per month during two hours prior to sunrise. Ψ_PD_ was measured using a Scholander pressure chamber (Model 3000 Plant Water Status Console, Soil Moisture Equipment Corp., Santa Barbara, USA). Chlorophyll fluorescence, specifically, maximum photochemical efficiency of photosystem II (PSII) of dark-adapted leaves (Fv/Fm), was assessed using a MINI-PAM-II (Heinz Walz GmbH, Effeltrich, Germany).

#### On-line gas exchange measurements

An automated measurement system was installed inside the climate chambers enabling the online assessment of leaf gas exchange (Werner et al. 2020; Meischner et al. 2024). Each of the two climate chambers was equipped with 14 connections for cuvettes. A defined gas flow of purified air (zero-air generator, ZA-FID-AIR, LNI Swissgas GmbH, Kamen, Germany) was directed through the cuvettes and subsequently analysed for H_2_O, CO_2_, and ^2^H by an analyser system. One branch per plant was enclosed in each cuvette, which were made of borosilicate glass (780 mL volume). Cuvettes were closed with Nalophan foil, sealed air tight and equipped with a fan (MC25101V2-000U-A99, Sunon, Kaohsiung City, Taiwan) to ensure homogeneous gas mixture and to prevent condensation. The air inflow into the cuvettes was controlled by mass flow controllers at a constant rate of 400 ml min^-1^. The air flow out of the cuvettes was regulated at 250 ml min^-1^, resulting in a slight overpressure inside the cuvettes to avoid contamination of measured air. The outflow of the cuvettes was directed through gas multi position valves (14 positions, 1/8”, Valco Instruments CO. Inc., Schenken, Switzerland) enabling switching between the cuvettes, measuring one after another for six minutes each. The analyser system consisted of an infrared gas analyser (LI-850, LI-COR Environmental, Bad Homburg, Germany) for CO_2_ and H_2_O concentration measurements and a water isotope laser spectroscope (L2130-i, Picarro Inc., Santa Clara, USA) analysing the H_2_O vapour concentration and δ^2^H isotopic signature. Two reference air cuvettes were left empty to determine gas concentrations of the air flowing into the cuvettes. Leaf and projected needle area of the measured branches were determined non-destructively, based on pictures, using the open access application Easy Leaf Area (Easlon and Bloom 2014). The concentration differences between the reference air and the leaf cuvettes were used to calculate net CO_2_ assimilation (*A,* µmol m^-2^ s^-1^) and transpiration rates (*E,* mmol m^-2^ s^-1^), and stomatal conductance (*g_s_*, mmol m^-2^ s^-1^) was subsequently derived from *E* and leaf-to-air water vapour mole fraction difference (ΔW), all according to von Caemmerer and Farquhar (1981).

#### VOC emission measurements

For the VOC sampling, glass thermodesorption tubes (Gerstel, Müllheim a. d. R., Germany) filled with Tenax TA (Sigma Aldrich, Munich, Germany) were connected to the outlet of each cuvette and air sampling pumps (Pocket Pump TOUCH, SKC Inc., Pittsburgh, USA) for two hours at a flow rate of 100 mL min^-1^. The thermodesorption tubes were analysed by gas chromatography-mass spectrometry (GC-MS 7890B GC System & 5975C MSD, Agilent Technology, Santa Clara, USA) according to (Haberstroh et al. 2018; Kreuzwieser et al. 2021). The Agilent Mass Hunter software (Agilent Technologies, Böblingen, Germany) was used for compound identification with the NIST mass spectral library and peak area quantification. Peak alignment and, if necessary, correction of the peak area integration was performed manually. A standard mixture of the following compounds was used for the quantification: α-pinene, β-pinene, limonene, trans-β-ocimene and trans-β-caryophyllene. For the quantification of monoterpenes (MTs) and sesquiterpenes (SQTs) not covered by the standard mixture, α-pinene and trans-β-caryophyllene were used, respectively. In total, nine compounds could be detected for *F. sylvatica*, 21 for *P. menziesii* and 27 for *P. abies* (Supplementary Material, Table S1).

#### Water-soluble carbohydrates

Water-soluble organic matter (WSOM) was determined based on the anthrone method according to Yemm and Willis (1954). Approximately 30 mg of dried leaf material was extracted in 1.0 mL distilled water at 90 °C for 5 min. Homogenates were centrifuged at 14,000 × g for 5 min, and the supernatant was diluted 1:10 with distilled water. For the colorimetric assay, 200 µL of the diluted extract was mixed with 1 mL anthrone reagent (0.5 g L^-1^ anthrone, 10 g L^-1^ thiourea in 70% H₂SO₄). Samples were incubated in a boiling water bath for 15 min, cooled on ice, and absorbance was measured at 578 nm using a spectrophotometer (DU 650, Beckman Coulter Life Sciences, Indianapolis, USA). Carbohydrate concentrations were calculated based on a sucrose standard curve (0–100 nmol) and expressed as mg per g dry weight (mg g^-1^).

### Statistical analysis

Data processing and statistical analyses were performed using the open-source program R (version 4.5.1, R Core Team, 2025). Significance levels of p < 0.05 (*), p < 0.01 (**), and p < 0.001 (***) were applied throughout.

To compare physiological parameters between mortality groups or species, normality of the data was evaluated visually and statistically with the Shapiro-Wilk test. When the assumption of normality was met and sample sizes were sufficient, Welch’s t-test was used, otherwise the non-parametric Wilcoxon-Test was employed (both implemented in the *rstatix* package (Kassambara 2025)). P-values were adjusted for multiple comparisons using the Bonferroni correction.

Diurnal δ^2^H values were averaged per plant and day and subsequently per mortality group and species to calculate means ± standard error (SE). Temporal trends were fitted using a four-parameter logistic sigmoid model using the *SSfpl* function from base R. The validity of the model fit was verified by testing the normality of residuals visually.

## Results

### Meteorological conditions

2023 was the warmest year on record to date in Germany, with exceptionally high T_air_ even during autumn (German Meteorological Service, DWD, 2023). The rain-out shelter additionally increased T_air_ compared to ambient conditions by 0.87 ± 0.71 °C (range: -0.82 to 4.31 °C), particularly at days with high solar radiation (Fig. S1). From 06^th^ of July until 13^th^ of October, a total of 52 hot days were recorded, with the latest hot day occurring on 13^th^ of October. Most of the hot days (73.1%) were observed during July and August (Fig. 2a,d). Highest recorded T_air_ under the rain-out shelter was 39.3 °C (9^th^ of July 2023). Moreover, we observed a total of eight tropical nights, six of them occurring during August (Fig. 2a,d). VPD reached maximum values of 4.6 kPa (Figure 2b). PPFD was above 1000 µmol m^-2^ s^-1^ on 65 out of 118 measured days (Fig. S2). Regular watering maintained volumetric water content (VWC) close to the targeted range of 20–30 % throughout the growing season to avoid edaphic drought, with the 5th to 95th percentile ranging from 19–31% (median: 25%) across all sensors during June – October 2023. A cold spell during the beginning of 2024 with 14 consecutive frost days (08^th^ – 21^st^ of January 2024), induced frost desiccation, indicated by a rapid decline of VWC in all boxes to a mean minimum of 13 ± 4 % across all boxes on 16^th^ of January 2024 (Fig. 2c,d).

**Fig. 2.**
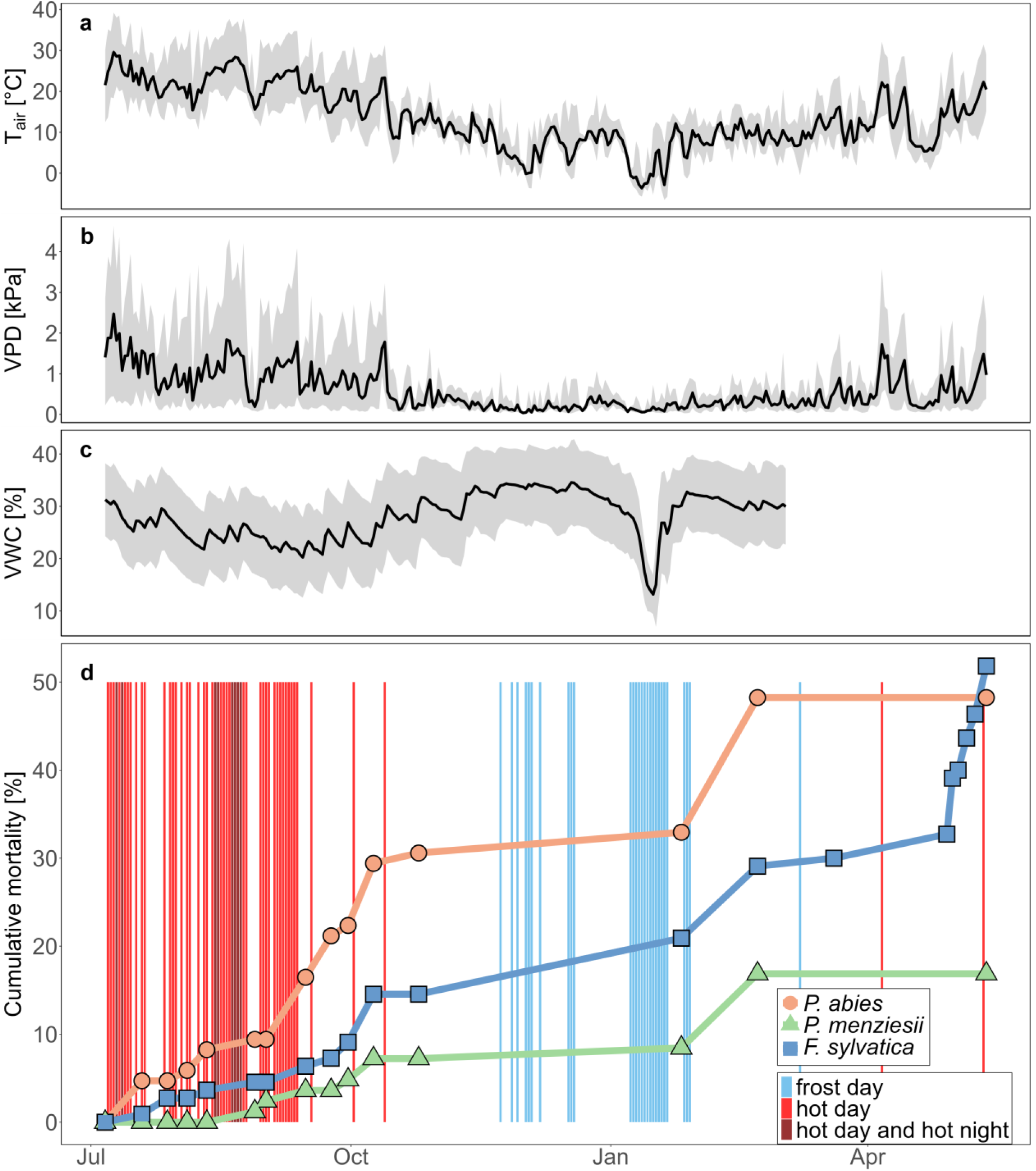
Daily mean air temperatures (Tair) (a) and vapor pressure deficit (VPD) (b) under the rainout-shelter with daily maximum and minimum values (shaded areas). (c) shows daily mean volumetric water content of the measured boxes with standard deviation (shaded area). (d) indicates the cumulative mortality [%] per species over the observed period. Vertical bars (d) indicate frost days (minimum daily T_air_ < 0 °C), hot days (maximum daily T_air_ > 30 °C) and hot days and hot nights under the rain-out shelter (minimum daily T_air_ > 20 °C).

### Tree mortality

Overall mortality, including plants that died during 2023 and during spring 2024, was highest in *F. sylvatica* (51.8%), followed by *P. abies* (48.2%) and *P. menziesii* (16.9%) (Fig. 3a). *F. sylvatica* and *P. menziesii* had higher mortality rates during spring 2024 than in 2023, in contrast to *P. abies*, where most of the saplings (30.6%) died in 2023 (Fig. 3a). Mortality in *P. abies* started with the onset of the heat waves in July 2023 and progressed rapidly, whereas the other two species showed increased mortality only towards the end of the season (Fig. 2d). Time between first discoloration of the leaves and final death was shortest in *P. abies,* which differed significantly from *P. menziesii* (p < 0.01) and *F. sylvatica* (p < 0.001) (Fig. 3b).

**Fig. 3.**
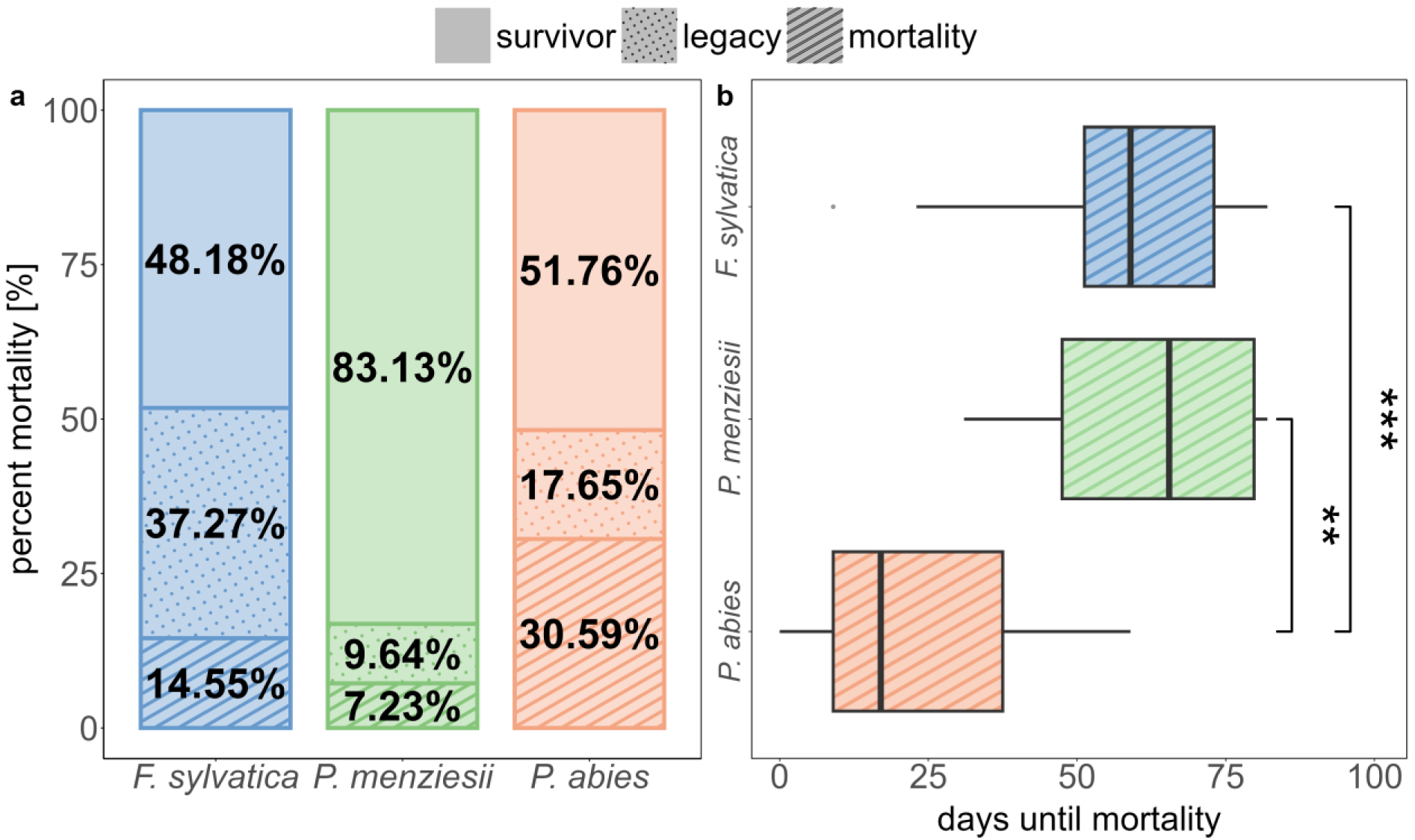
(a) Proportion of plants that died during season 2023 (mortality), died as legacy effect in beginning of 2024 (legacy) or survived (survivor) for *F. sylvatica* (n=110), *P. menziesii* (n=83) and *P. abies* (n=85). (b) number of days until mortality per species since first documentation of leaf discoloration for the respective individual. (b) only includes the saplings that died during vegetation season 2023 (mortality). Asterisks indicate significant differences between species (p < 0.01 = **, p < 0.001 = ***).

### Physiological responses

#### Water relations and water uptake velocity

Water relations during July–September exhibited clear differences between the groups with highest performance in the survivor population of each species and a clear gradient with increasing susceptibility to mortality (survivor > legacy > mortality), characterized by decreasing stomatal conductance (*g_s_*), transpiration rates (*E*) and predawn leaf water potential (Ψ_PD_) (Fig. 4). For instance, compared to surviving plants, *g_s_* in the legacy and mortality groups was reduced by 22.0 % and 67.8 % in *F. sylvatica*, by 58.1 % and 94.0 % in *P. menziesii*, and by 72.5 % and 75.2 % in *P. abies* during July–September. By October, differences in *E* and *g_s_* between the survivor and legacy groups of the conifers became even more pronounced (p < 0.001). At this point, *g_s_* in legacy plants was reduced by 83.3 % in *P. menziesii* and by 85.4 % in *P. abies* relative to survivors. In *F. sylvatica,* no significant differences emerged between the groups during October for none of the monitored parameters, but both groups exhibited reduced *E* and *g_s_* compared to the earlier measurements during July–September, probably due to the onset of leaf senescence. In all species, reduction of Ψ_PD_ in the legacy groups generally showed smaller deviations from Ψ_PD_ in survivors than observed for *E* and *g_s_*. Ψ_PD_ only differed significantly for *P. abies* between legacy and survivor group in October (Fig. 4). Plant water use and uptake velocity differed strongly between the species and across the mortality groups, as indicated by diverging ^2^H uptake dynamics (Fig. 5). From the δ^2^H increase following pulse labelling, we calculated the time of the maximum uptake rate defined as the inflection point (X_mid_) of the fitted logistic curve. Legacy plants of *F. sylvatica* and *P. abies* responded with a time delay of up to 1.6 days until the maximum uptake rate, and overall significantly (p < 0.001) later compared to the survivors. Differences in response time between the legacy and survivor group of *P. menziesii* were minor (0.16 days) compared to the other two species and non-significant (Fig. 5, Table 1). Maximum values of δ^2^H of transpired water ranged between 488.2 ± 234.9 and 645.3 ± 396.2 ‰ across all survivor groups. In the legacy plants of *F. sylvatica* and *P. menziesii*, they were significantly (p < 0.05) reduced by 72.5 % and 62.7 % compared to the survivor plants. Contrastingly, in *P. abies*, despite the delayed response, maximum δ^2^H values in legacy plants and mortality plants were only slightly, non-significantly reduced compared to survivor plants by 13.0 % and 10.3 %, respectively (Fig. 5, Table 1).

**Fig. 4.**
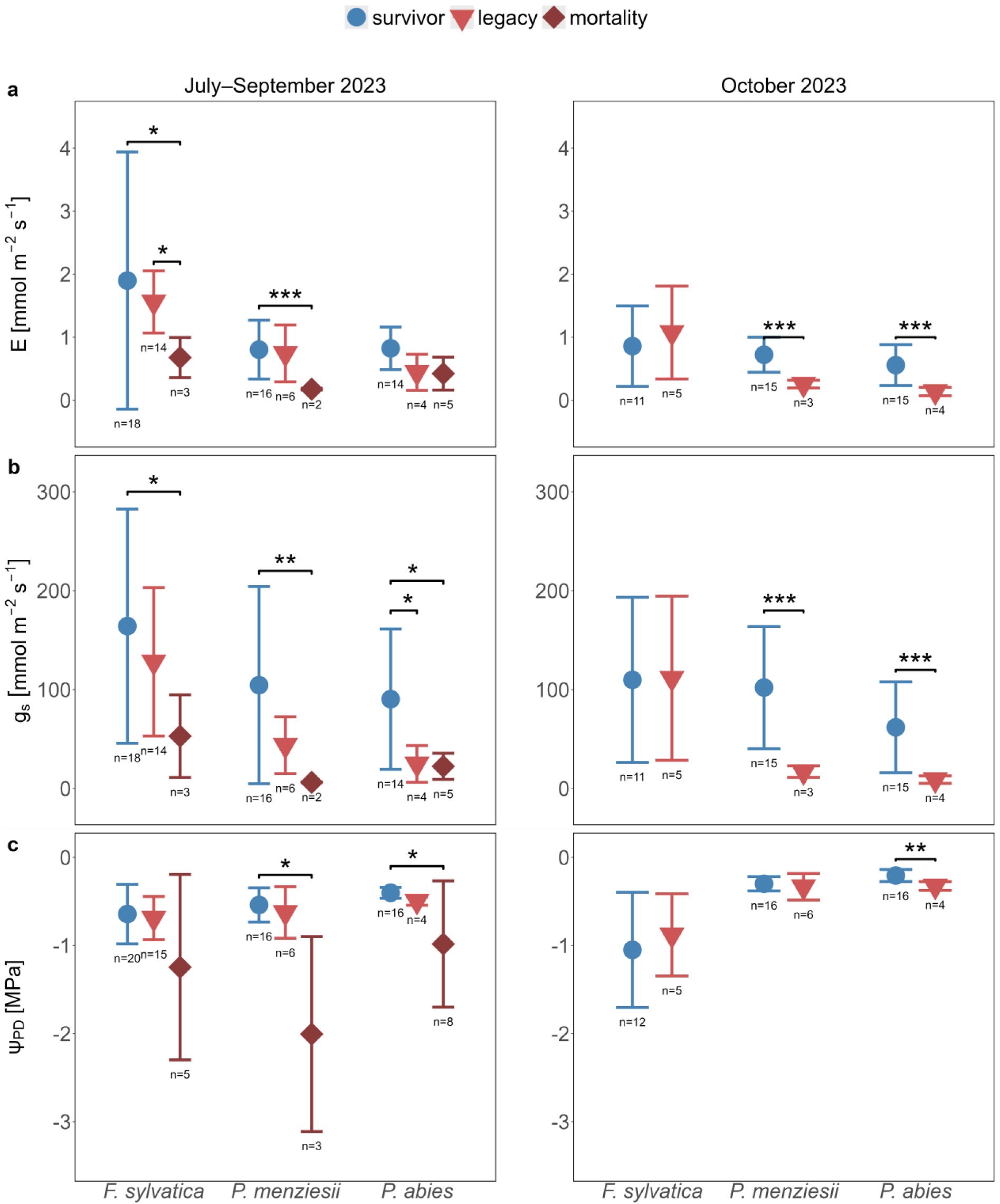
Water relations across all mortality groups during 2023 growing season (left panels) and survivor and for legacy plants in October 2023 (right panels). Dots show means ± standard deviation (SD) per species and mortality group. (a) Transpiration (*E*) and (b) stomatal conductance (*g_s_*) only include daytime values and were measured in climate chambers at 28 and 23 °C during July–September and October 2023, respectively. (c) Predawn leaf water potential (Ψ_PD_) was measured outside at ambient T_air_ prior to sunrise.

**Fig. 5.**
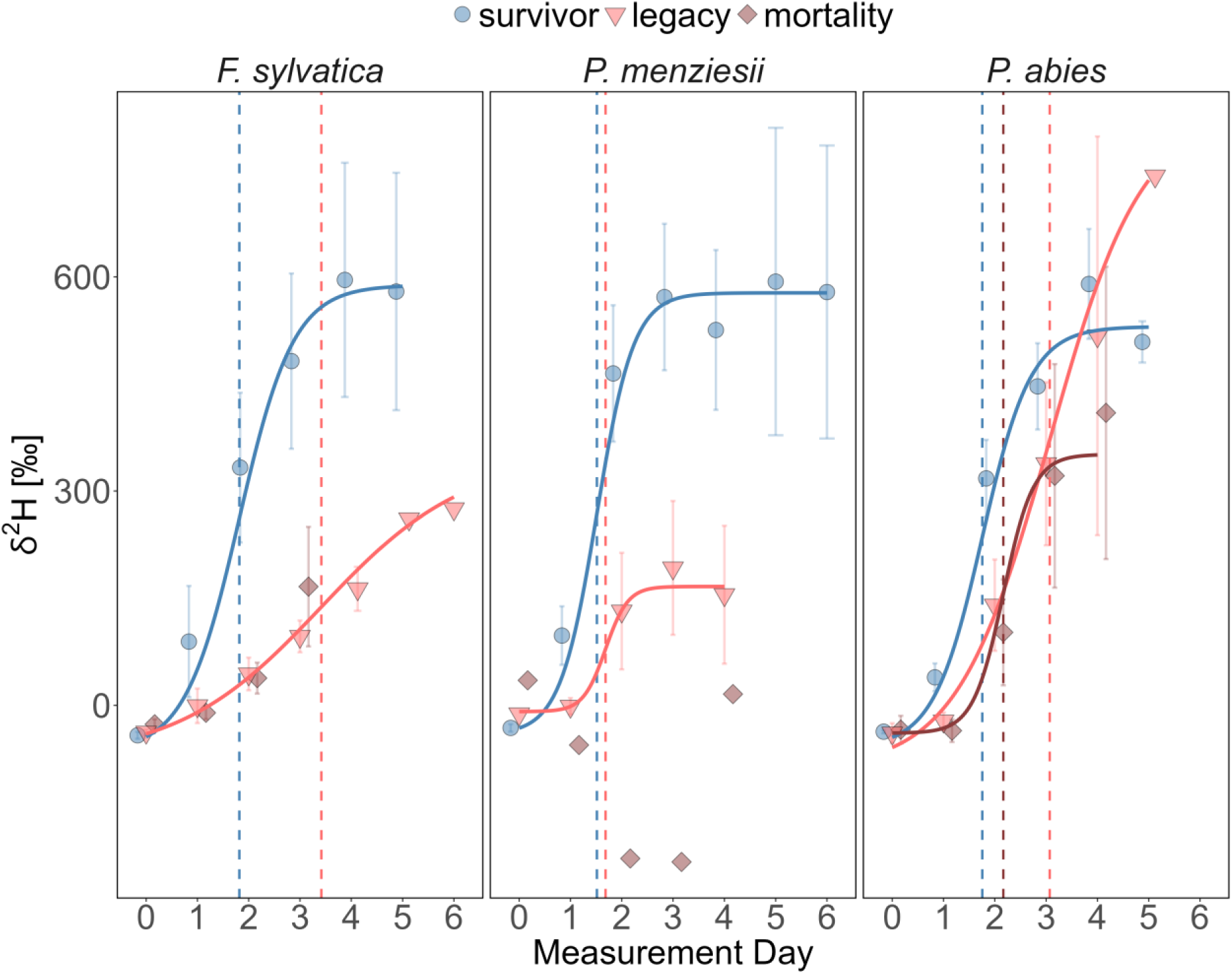
Temporal dynamics of hydrogen isotope signature (δ^2^H) in response to ^2^H-enriched irrigation. Symbols represent daily mean δ^2^H ± standard error (SE) per species and mortality group. Daily means were calculated from individual plant means, which were derived from daytime measurements only. Measurement day zero corresponds to baseline measurements taken during the first measurement period in the climate chambers (M1). Irrigation with ^2^H-enriched water was applied on measurement day 1. Solid lines show fitted four-parameter logistic sigmoid models for species-mortality groups with n ≥ 3. Vertical dashed lines indicate the inflection points (X_mid_) of the fitted models. All measurements were performed at T_air_ = 28 °C.

**Table 1:** Hydrogen isotope signature (δ²H) dynamics following ²H-enriched irrigation in *F. sylvatica*, *P. menziesii* and *P. abies* across mortality groups.

| Species | Group | n | $X_{\text{mid}}$ [day of experiment] | $\Delta^2\text{H}_{\text{max}}$ [‰] ± SD |
| --- | --- | --- | --- | --- |
| *F. sylvatica* | Survivor | 12 | 1.82<sup>a</sup> | 576 ± 504<sup>a</sup> |
|  | Legacy | 7 | **3.42<sup>b</sup>** | **158 ± 98<sup>b</sup>** |
| | Mortality | 2 | NA | 226 ± 194<sup>ab</sup> |
| *P. menziesii* | Survivor | 13 | 1.52<sup>a</sup> | 645 ± 396<sup>a</sup> |
|  | Legacy | 5 | 1.68<sup>a</sup> | **241 ± 180<sup>b</sup>** |
|  | Mortality | 1 | NA | 52 |
| *P. abies* | Survivor | 14 | 1.76<sup>a</sup> | 488 ± 235<sup>a</sup> |
| | Legacy | 4 | **3.10<sup>b</sup>** | 425 ± 270<sup>a</sup> |
| | Mortality | 3 | 2.16<sup>a</sup> | 438 ± 374<sup>a</sup> |

Number of replicates (n), maximum uptake velocity ( $X_{\text{mid}}$ ) and mean maximum $\delta^2\text{H} \pm$ standard deviation (SD) per mortality group and species.
Letters (a, b, ab) indicate significant differences between the mortality group within one species.
Groups with $n < 3$ were not tested for statistical differences due to low number of replicates.
Significantly higher values (in intraspecific comparison) are highlighted in bold ( $p < 0.05$ ).

### Regulation of carbon uptake and VOC emissions

Net assimilation (*A*) and predawn Fv/Fm generally followed the observed hierarchy in water dynamics (survivor > legacy > mortality), with differences between legacy and survivor groups becoming significant in October for both conifers (p < 0.01) (Fig. 6). Differences between the groups were more pronounced in *A* than in Fv/Fm, which generally showed weaker declines in the legacy group, similarly to Ψ_PD_. Mean Fv/Fm values in October were 0.79 ± 0.07 and 0.81 ± 0.02 in the survivor group, while the values in the legacy group had decreased to 0.67 ± 0.20 and 0.54 ± 0.25 in *P. menziesii* and *P. abies*, respectively, indicating persistent stress without the presence of drought or heat.

**Fig. 6.**
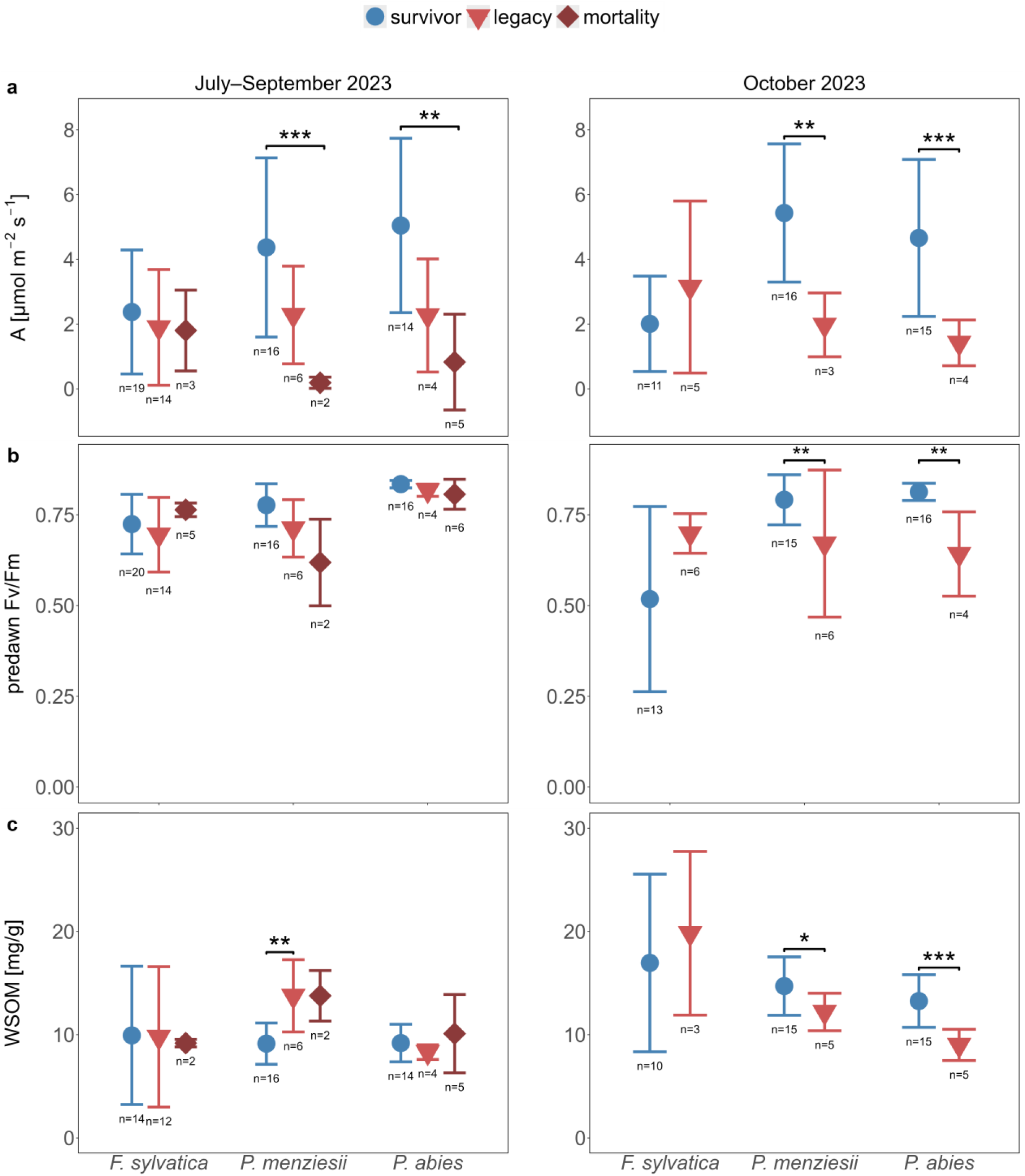
Carbon related measurements across all mortality groups during 2023 growing season (left panels) and for survivor and legacy plants in October 2023 (right panels). Dots show means ± SD per species and mortality group. (a) Net assimilation (*A*) and (c) WSOM were measured in climate chambers at 28 and 23 °C during July–September and October 2023, respectively. *A* only includes daytime values. (b) Predawn ChlF was measured outside at ambient T_air_ prior to sunrise.

In *P. menziesii*, concentration of WSOM in leaf tissue initially was higher during July–September in legacy (p < 0.01) and mortality plants (non-significant) compared to the survivor plants, while in *F. sylvatica* and *P. abies* variations between the mortality groups were minor. During October, WSOM concentrations increased in the survivor groups of all three species and in the legacy group of *F. sylvatica*, resulting in significantly higher concentrations in the survivor group of *P. menziesii* (p < 0.05) and *P. abies* (p < 0.001) compared to the legacy groups (Fig. 6).

During July–September, survivor and legacy plants exhibited comparable total emissions across all species, varying only slightly in composition (Fig. 7a). However, mortality plants differed strongly from legacy and survivors in all species. Total VOC emissions in mortality plants of *F. sylvatica* and *P. abies* increased by 147.8% and 66.6%, respectively, compared to the survivors, whereas mortality plants of *P. menziesii* decreased total emissions by 90.2% compared to survivors (Fig. 7). The greatest emission increase in mortality plants during this period was observed for the monoterpene sabinene. In *F. sylvatica*, sabinene emissions increased by 397.2 and 464.0 nmol m^-2^ h^-1^ relative to survivor and legacy plants, respectively. In *P. abies*, the increase was 104.1 and 107.7 nmol m^-2^ h^-1^. Mortality plants of *F. sylvatica* further showed higher emission rates of the monoterpene cyclofenchene (+60.7 and +62.7 nmol m^-2^ h^-1^), while mortality plants of *P. abies* emitted the monoterpenes camphene (+69.1 and +72.7 nmol m^-2^ h^-1^) and β-myrcene (+80.3 and +83.8 nmol m^-2^ h^-1^) and the aldehyde nonanal (+34.1 and +34.7 nmol m^-2^ h^-1^) at higher rates. In *P. menziesii*, greatest compositional differences were detected for the alcohol 2-propen-1-ol and the monoterpene β-pinene, which were emitted by survivor and legacy plants but were absent in mortality plants. However, it has to be acknowledged that the mortality group only consisted of two individuals.

**Fig. 7.**
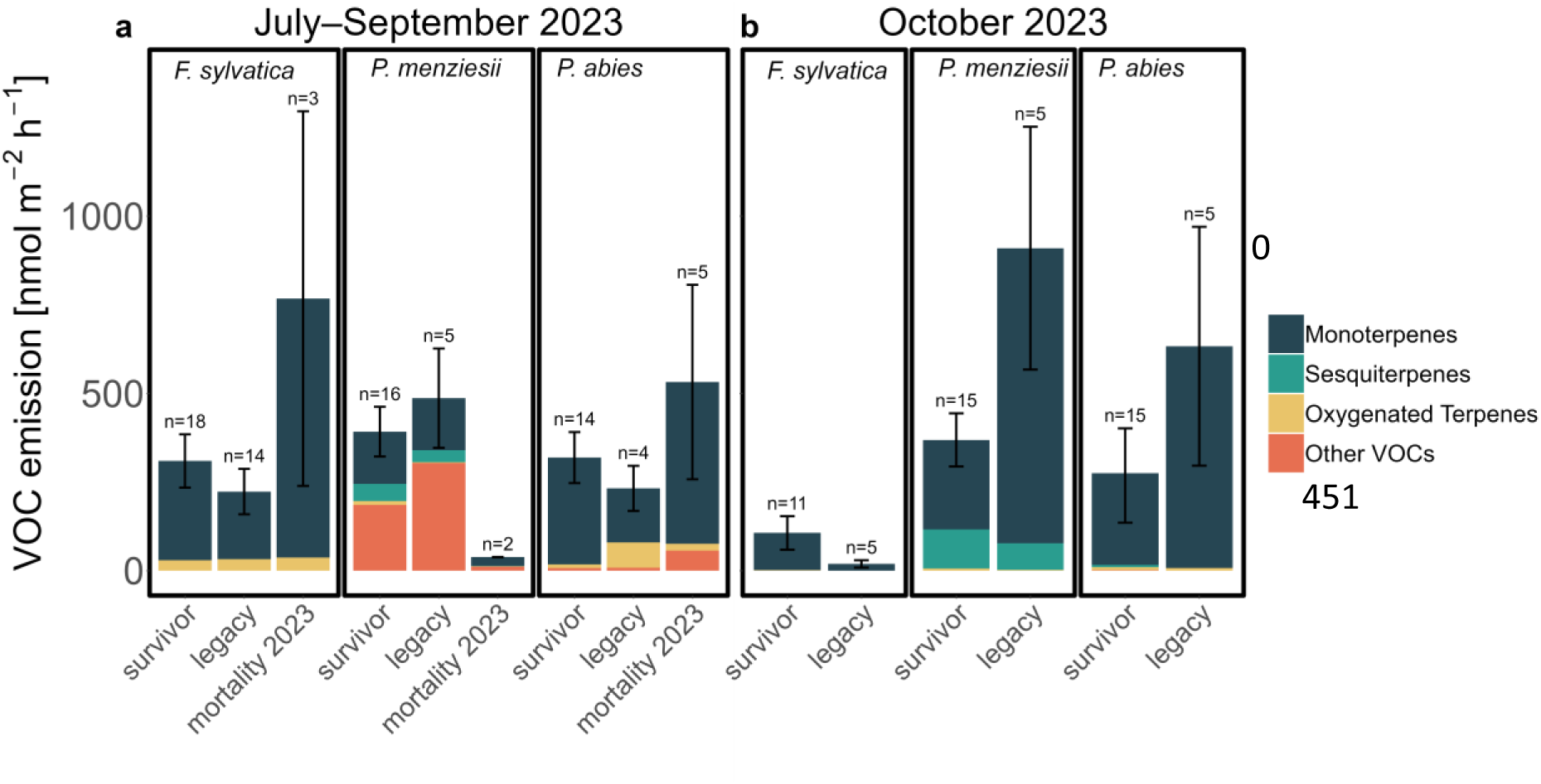
VOC emissions during July–September 2023 (a) and during October 2023 (b) with standard error for total emission. Data include all identified VOC emissions for each mortality group of the three species.

Importantly, in October 2023, legacy plants of both conifers exceeded total emissions of the survivor plants due to higher monoterpene emissions (Fig. 7b). During October 2023 and relative to survivors, total emissions of the legacy plants increased by 146.5% and 135.6% in *P. menziesii* and *P. abies*, respectively. In *P. menziesii*, this was driven primarily by higher emissions of monoterpenes β-pinene (+400.2 nmol m^-2^ h^-1^) and α-pinene (+184.9 nmol m^-2^ h^-1^) in the legacy individuals, while legacy plants of *P. abies* mainly increased monoterpenes β-phellandrene (+330.9 nmol m^-2^ h^-1^) and δ^3^-Carene (+56.2 nmol m^-2^ h^-1^). In contrast, *F. sylvatica* showed reduced total emissions in October 2023 compared to July–September in both groups, with the survivor plants maintaining higher rates than the legacy plants (Fig. 7). This was mainly caused by higher emissions of sabinene (+33.4 nmol m^-2^ h^-1^), β-phellandrene (+29.2 nmol m^-2^ h^-1^) and δ^3^-carene (+20.7 nmol m^-2^ h^-1^) in the surviving plants relative to the legacy plants.

## Discussion

Globally, forest ecosystems are experiencing widespread mortality, yet predicting dieback events and identifying their drivers remains a critical challenge (Allen et al. 2015; Hartmann et al. 2022; Werner et al. 2026). This is compounded by the fact that lethal physiological thresholds in trees are typically exceeded long before visual symptoms appear (Blackman et al. 2019; Hammond et al. 2019), limiting opportunities for intervention. Consequently, future forest management requires physiological warning indicators, enabling interventions that mitigate ecological and economic damages (Mosig et al. 2026). Here, we demonstrate that heat stress alone - independent of drought - can drive mortality and increase vulnerability to subsequent stress events in temperate tree species. We identify reduced water uptake velocity and stomatal conductance, impaired photosynthetic efficiency, and altered terpenoid emissions as early warning indicators, distinguishing survivors from plants that eventually succumbed to legacy effects months before visible damage occurred. These findings are highly useful for establishing a physiological framework for predicting tree mortality and provide a foundation for developing monitoring protocols and improving mortality models in the context of climate change.

### Heat waves threaten temperate forest vitality

During recent decades, Central European temperate forests have experienced unprecedented T_air_ extremes and elevated canopy temperatures, yet their acclimation capacity remains unclear (Werner et al. 2026). Recent mortality has particularly affected *P. abies* and *F. sylvatica* (Schuldt et al. 2020), two of the most abundant temperate tree species in Germany (Obladen et al. 2021; Riedel et al. 2024). While primarily attributed to drought or compound drought (Schuldt et al. 2020; Senf et al. 2020), increasing evidence indicates heat and elevated VPD can operate as independent drivers of mortality (Birami et al. 2018; Grossiord et al. 2020; Still et al. 2023; Werner et al. 2026). Our results confirm high vulnerability of *F. sylvatica* and *P. abies* saplings to heat exposure. Summer heat induced severe mortality rates in well-watered *F. sylvatica* and *P. abies* and subsequent winter desiccation triggered further mortality. In contrast, *P. menziesii* exhibited substantially lower mortality, indicating greater heat resilience. As morphological adaptations (e.g. deep rooting) cannot explain these patterns in potted saplings, our results point to inherently higher hydraulic safety in *P. menziesii*, likely mediated by higher wood density and cuticle properties being present from the sapling stage (Dalla-Salda et al. 2009; Leuschner and Meinzer 2024). Mortality screenings revealed significantly faster progression from initial to complete needle discoloration and a higher proportion of immediate mortality in *P. abies* compared to the other species (Fig. 3), suggesting rapid hydraulic failure in *P. abies.* This is in agreement with observations in mature trees following natural drought (Arend et al. 2021), aligning with its identification as Europe’s most vulnerable conifer (Spiecker et al. 2019). Contrastingly, damage in *F. sylvatica* and *P. menziesii* progressed more gradually. Thus, delayed mortality in *F. sylvatica* and *P. menziesii* may enable early intervention if physiological indicators are detected before lethal thresholds are crossed.

### Heat exposure impairs hydraulic function in legacy and mortality plants

Heat exposure and elevated VPD increase atmospheric evaporative demand (Grossiord et al. 2020; Novick et al. 2024; Werner et al. 2026). An imbalance between tree water supply and demand can impair xylem conductivity, potentially causing hydraulic failure and whole tree mortality (Anderegg et al. 2013; Adams et al. 2017; Werner et al. 2026), which has often been shown in the context of combined edaphic and atmospheric drought (Birami et al. 2018; Arend et al. 2021). For instance, adult *F. sylvatica* and *P. abies* trees, which were previously drought stressed, exhibited slower label water uptake after ^2^H labelling compared to control trees, indicating damages to the hydraulic system (Hesse et al. 2026). Hesse et al. (2026) proposed changes in soil and root properties, impaired water transport, or the replenishment of water reserves as possible explanations. In our study, despite identical treatments, legacy and mortality individuals across all species exhibited reduced water uptake velocities compared to the survivors after heat exposure (Fig. 5). This expands previous findings by implying a loss of hydraulic conductivity caused by heat stress alone. The reduced water uptake velocity in the legacy and mortality plants likely reflected early heat-induced hydraulic damages, which may be attributed to differing stomatal behaviour, structural damage of stomata and lower protective capacities of the respective individuals compared to survivors. These findings highlight the need for individual-tree monitoring approaches that capture intraspecific variation in vulnerability to environmental stress.

Reduced water uptake velocities further influence carbon allocation dynamics (Rehschuh et al. 2022). In survivor plants, WSOM dynamics followed established seasonal patterns, decreasing during the growing season and replenishing towards dormancy (Furze et al. 2019; Acevedo-Siaca et al. 2021). However, legacy and mortality plants, particularly in the conifers, deviated from this pattern. Elevated WSOM concentrations in legacy and mortality plants of both conifers during the growing season likely reflect impaired transport capacity. In contrast, survivor conifers maintained carbon allocation capacity, exhibiting depleted WSOM concentrations during the growing season and replenishing reserves prior to winter dormancy (Fig. 6) (McDowell et al. 2008; Anderegg et al. 2012).

### Conservative stomatal behaviour increases vulnerability to mortality during heat waves

We identified consistent intraspecific variation in stomatal behaviour across all species following heat stress, with stomatal conductance following a clear gradient (survivor > legacy > mortality) despite comparable soil water availability. Reduced stomatal conductance in legacy and mortality plants likely reflected reduced water uptake capacity and/or heat-induced tissue damages. However, Ψ_PD_ values indicate legacy plants replenished water reservoirs over night during the 2023 growing season, whereas mortality plants showed significantly reduced Ψ_PD_ (Fig. 4).

Plants typically close stomata to prevent water loss. During heat exposure, this creates a critical trade-off between transpirational cooling and water conservation (Bachofen et al. 2025; Kullberg et al. 2026). Closed stomata restrict transpirational cooling, causing leaf temperatures to rise (Cochard 2021). Exceeding species-specific leaf temperature thresholds damages the photosynthetic system, e.g. PSII (Hüve et al. 2011; Guha et al. 2018), and increases cuticular conductance, potentially inducing sudden run-away embolism and mortality (Arend et al. 2021; Cochard 2021; Garen and Michaletz 2025). To avoid such thermal damages, isohydric plants have even shown to shift to anisohydric behaviour under heat (Sharma et al. 2026). In our study, decreased Fv/Fm values during October 2023 in the legacy plants of the two conifers - without the presence of environmental stress - (Fig. 6) indicate damages to the photosynthetic apparatus, likely triggered by increased leaf temperatures resulting from reduced stomatal conductance. Most legacy conifers died shortly after winter desiccation, suggesting reduced water supply from frozen soil, combined with pre-existing damage, failed to meet water demand, resulting in hydraulic failure.

These patterns suggest that individuals with conservative stomatal behaviour and lower transpiration were inherently more vulnerable to heat-induced mortality and legacy effects across all species, consistent with observations that transpirational cooling capacity increases the likelihood of survival during heat stress (Kolb and Robberecht 1996; Birami et al. 2018). Accordingly, model simulations confirm that plant survival during heatwaves critically depends on stomatal state at exposure, with heatwaves following stomatal closure provoking total xylem embolization within hours, whereas open-stomata conditions during heat waves had minor mortality impacts (Cochard 2021). Intraspecific differences in hydraulic management likely reflect both reduced heat resilience and genetic variability expressed through stomatal density, size, or behaviour (McDowell et al. 2008). Critically, our results demonstrate that reduced stomatal conductance exacerbates heat effects even under well-watered conditions, with early signals occurring up to 10 months before mortality.

### VOC emissions reveal thermal stress long before mortality occurs

Heat exposure significantly alters VOC emissions by increasing their volatility and by enhancing the production of specific VOCs with thermoprotective capacities, such as stabilizing chloroplast membranes and scavenging reactive oxygen species (Peñuelas and Llusià 2003; Vickers et al. 2009; Loreto and Schnitzler 2010). VOC emissions can therefore serve as valuable indicators of heat-induced stress and reveal whether and to which extent plants allocate carbon towards protective mechanisms (Werner et al. 2020). Additionally, a previous study has found altered VOC emissions in dying individuals of *Pinus halepensis* saplings before gas exchange parameters reacted and up to 14 days before mortality occurred (Birami et al. 2021), which highlights the further potential of VOC measurements as early indicators for mortality. Our results confirm this, as we observed increased VOC emission in *F. sylvatica* and *P. abies* plants shortly before they died, suggesting the activation of protective responses against heat-induced damage. Elevated emissions of specific monoterpenes in mortality plants of *F. sylvatica* (e.g. sabinene, cyclofenchene) and *P. abies* (e.g. sabinene, camphene, β-myrcene) might indicate targeted protective responses to the higher degree of thermal damage in these plants compared to the less affected *P. menziesii* saplings. Moreover, the increased emission of the aldehyde nonanal in *P. abies* could also be related to passive leakage resulting from structural damage by oxidative stress (Wildt et al. 2003; Loreto et al. 2006; Lee et al. 2026). However, in addition to the altered emissions in plants that died shortly after, we could also detect increased monoterpene emissions in legacy plants of both conifers before dormancy (Fig. 7). Notably, the total emission increases of 146.5% and 135.6% in legacy plants of *P. menziesii* and *P. abies*, respectively, occurred four months before they succumbed to legacy effects. These findings even expand the potential of VOC emissions as indicator for mortality, as emission changes occurred significantly earlier as previously observed. Such findings do not only indicate heat-induced damages, but also highlight the vulnerability of individuals to future stressors and might therefore serve as valuable parameters for forest management or seedling selection to mitigate the effects of future stress events.

## Conclusions

With this study we demonstrate the high susceptibility to heat exposure in saplings of *F. sylvatica* and *P. abies,* resulting in severe immediate and delayed mortality, while *P. menziesii* exhibited higher resilience. We show that heat exposure influenced various physiological parameters, resulting in impairments of the hydraulic and the photosynthetic systems. Consequently, we demonstrate that heat events alone are sufficient to drive tree mortality, independent of concurrent edaphic drought. Crucially, we identify water uptake velocity, stomatal conductance, photosynthetic efficiency and VOC emissions as specific physiological indicators that signal individual vulnerability and sub-lethal damage long before visible changes occur. Our findings identify a range of physiological early warning indicators that could i) guide sapling selection for climate-resilient genotypes by identifying traits associated with vulnerability, and ii) support forest management by assessing sub-lethal damage after stress exposure, allowing for interventions such as mitigating subsequent stressors.

## Acknowledgements

We gratefully acknowledge financial support by the German Research Foundation via the CRC1537 (ECOSENSE, Project ID: 459819582) and the RU Forest Floor (WE 2681/13-1), and by the Studienstiftung des deutschen Volkes. We thank Alexandra Paul and Anne-Marie Schiphorst for assistance with IRMS analysis, Monika Eiblmeier for assistance with GC-IRMS analysis and Eva Schottmüller, Fabio Scarpa, Finn Reimold, Lennart Nettler, Phyllis Lua-Mellmann and Helen Vogt for support with the plant material and the measurements.

## Author contributions

CW and SH designed the study. CS, SD, MM, MW and KK performed the experiments and collected data. CS, SD, MM, JK and SH analysed data. CS, SD, MM, SH and CW interpreted data. CS, SD and SH wrote the manuscript with inputs from all authors. All authors critically reviewed the manuscript.

## Data Availability

The data that support the findings of this study are available from the corresponding author upon reasonable request

