## Supplement for "Early warning indicators for heat-induced mortality in temperate tree saplings"

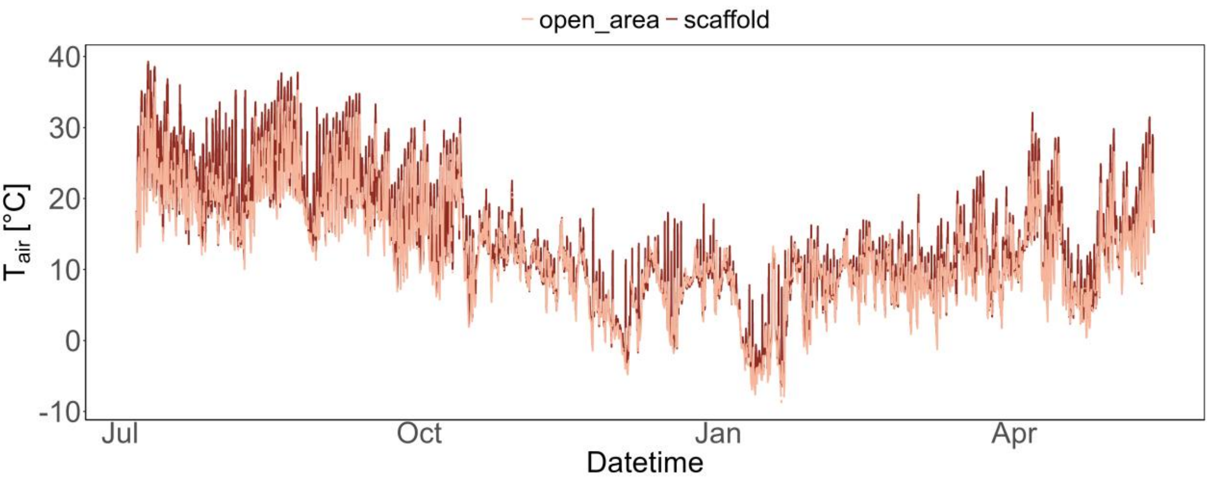


*Figure S1: $T_{air}$ measured every 5 min under the rain-out shelter (dark colour) and hourly at an open area nearby (bright colour).*

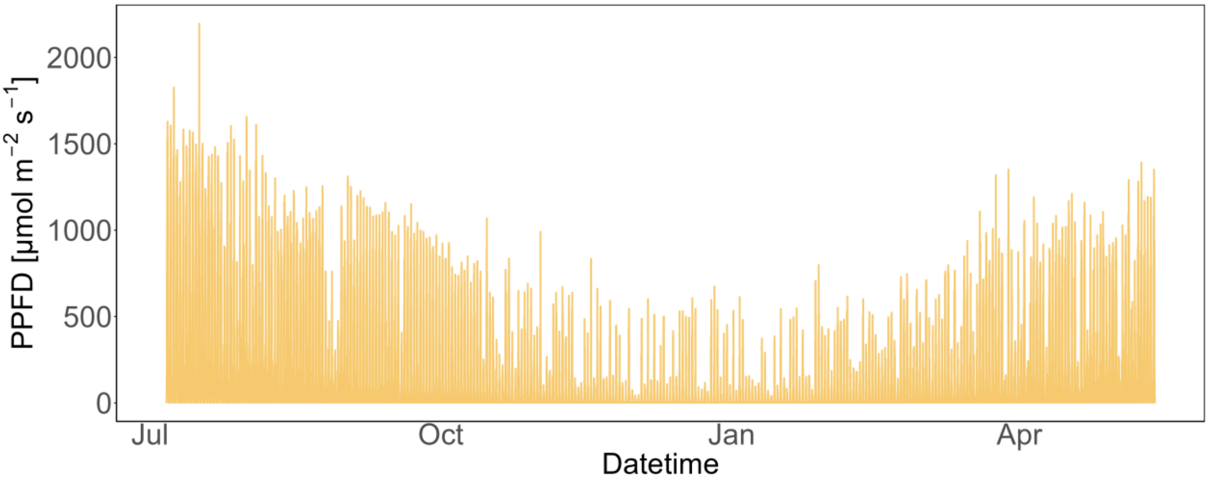


*Figure S2: PPFD under the scaffold, recorded every 5 min.*

Table S1: Identified VOCs measured by the GC-MS analysis per species.

| Compound | Species | Match Factor | CAS | Retention Time |
|---|---|---|---|---|
| α-Phellandrene | *F. sylvatica* | 85.52 | 99-83-2 | 25.67 |
| $\delta^3$-Carene | *F. sylvatica* | 94.46 | 13466-78-9 | 26.13 |
| Camphene | *F. sylvatica* | 87.36 | 79-92-5 | 27.15 |
| Sabinene | *F. sylvatica* | 97.71 | 3387-41-5 | 28.36 |
| β-Phellandrene | *F. sylvatica* | 89.04 | 555-10-2 | 28.69 |
| β-Myrcene | *F. sylvatica* | 93.37 | 123-35-3 | 29.08 |
| Cyclofenchene | *F. sylvatica* | 88.67 | 488-97-1 | 31.44 |
| γ-Terpinene | *F. sylvatica* | 86.43 | 99-85-4 | 32.75 |
| Geraniol | *F. sylvatica* | 89.41 | 124787-21-9 | 38.87 |

| | | | | |
|---|---|---|---|---|
| 2-Propen-1-ol | *P. menziesii* | 96.24 | 107-18-6 | 21.7 |
| Camphene | *P. menziesii* | 91.24 | 79-92-5 | 27.14 |
| β-Phellandrene | *P. menziesii* | 77.24 | 555-10-2 | 28.34 |
| β-Pinene | *P. menziesii* | 93.59 | 18172-67-3 | 28.69 |
| β-Myrcene | *P. menziesii* | 94.49 | 123-35-3 | 29.03 |
| α-Terpinene | *P. menziesii* | 91.03 | 99-86-5 | 29.74 |
| α-Pinene | *P. menziesii* | 97.48 | 80-56-8 | 30.26 |
| γ-Terpinyl acetate | *P. menziesii* | 90.66 | 10235-63-9 | 30.72 |
| p-Cymene | *P. menziesii* | 95.65 | 99-87-6 | 31.09 |
| Limonene | *P. menziesii* | 98.36 | 138-86-3 | 31.35 |
| 1,8-Cineole | *P. menziesii* | 90.06 | 470-82-6 | 31.54 |
| $\delta^3$-Carene | *P. menziesii* | 91.87 | 13466-78-9 | 32 |
| γ-Terpinene | *P. menziesii* | 86.94 | 99-85-4 | 32.69 |
| α-Thujone | *P. menziesii* | 81.15 | 546-80-5 | 35.02 |
| 1,2-Dimethoxybenzene | *P. menziesii* | 75.9 | 91-16-7 | 35.75 |
| p-Mentha-1,2-dien-8-ol | *P. menziesii* | 62.47 | 65293-09-6 | 36.48 |
| 1-Decanol | *P. menziesii* | 85.51 | 112-30-1 | 38.37 |
| Verbenone | *P. menziesii* | 96.35 | 80-57-9 | 38.74 |
| Geraniol | *P. menziesii* | 79.57 | 124787-21-9 | 38.87 |
| α-Bergamotene | *P. menziesii* | 80.4 | 13474-59-4 | 43.75 |
| α-Farnesene | *P. menziesii* | 93.42 | 502-61-4 | 44.75 |
| 2-Propen-1-ol | *P. abies* | 96.24 | 107-18-6 | 21.7 |
| $\delta^3$-Carene | *P. abies* | 94.02 | 13466-78-9 | 26.14 |
| Camphene | *P. abies* | 91.24 | 79-92-5 | 27.14 |
| Sabinene | *P. abies* | 88.44 | 3387-41-5 | 28.36 |
| β-Phellandrene | *P. abies* | 90.3 | 555-10-2 | 28.69 |
| β-Pinene | *P. abies* | 93.59 | 18172-67-3 | 28.69 |
| β-Myrcene | *P. abies* | 95.17 | 123-35-3 | 29.09 |
| α-Terpinene | *P. abies* | 91.03 | 99-86-5 | 29.74 |
| α-Pinene | *P. abies* | 97.48 | 80-56-8 | 30.26 |
| γ-Terpinyl acetate | *P. abies* | 90.66 | 10235-63-9 | 30.72 |
| β-Cyclocitral | *P. abies* | 73.81 | 432-25-7 | 30.79 |
| p-Cymene | *P. abies* | 95.65 | 99-87-6 | 31.09 |
| Limonene | *P. abies* | 98.36 | 138-86-3 | 31.35 |
| 1,8-Cineole | *P. abies* | 90.06 | 470-82-6 | 31.54 |

| | | | | |
|---|---|---|---|---|
| γ-Terpinene | *P. abies* | 86.94 | 99-85-4 | 32.69 |
| Nonanal | *P. abies* | 89.51 | 124-19-6 | 34.75 |
| α-Thujone | *P. abies* | 81.15 | 546-80-5 | 35.02 |
| 1,2-Dimethoxybenzene | *P. abies* | 75.9 | 91-16-7 | 35.75 |
| p-Mentha-1,2-dien-8-ol | *P. abies* | 62.47 | 65293-09-6 | 36.48 |
| Camphor | *P. abies* | 78.71 | 76-22-2 | 36.72 |
| α-Terpineol | *P. abies* | 76.37 | 2438-12-2 | 38.3 |
| 1-Decanol | *P. abies* | 85.51 | 112-30-1 | 38.37 |
| Verbenone | *P. abies* | 96.35 | 80-57-9 | 38.74 |
| Geraniol | *P. abies* | 79.57 | 124787-21-9 | 38.87 |
| α-Terpinyl acetate | *P. abies* | 84.17 | 80-26-2 | 42.02 |
| α-Bergamotene | *P. abies* | 80.4 | 13474-59-4 | 43.75 |
| α-Farnesene | *P. abies* | 93.42 | 502-61-4 | 44.75 |
